# Developing a Stepwise Induction Protocol for Cisplatin-Resistant Human Cervical Cancer Cells

**DOI:** 10.1101/2025.11.20.687683

**Authors:** Varsha Vijayakumar

## Abstract

Cisplatin, a platinum-based DNA crosslinking chemotherapeutic, remains the cornerstone of first-line chemoradiation therapy for advanced cervical cancer. However, the clinical efficacy of cisplatin is frequently compromised by the emergence of acquired resistance, which represents a principal cause of treatment failure and disease recurrence. Traditional protocols for generating resistant cervical cancer cell lines vary widely in dose-escalation schedules, passage criteria, timing, and endpoints, often requiring months to yield inconsistent results. To overcome these limitations, this study establishes a structured, stepwise protocol for developing cisplatin-resistant cervical cancer cell lines using HeLa (HPV18^**+**^) and SiHa (HPV16^**+**^) models. We hypothesised that a predefined dosing and monitoring scheme would shorten induction time while improving the reproducibility of resistance generation. Using this approach, we successfully generated cisplatin-resistant sublines (H_**1**_CR and S_**1**_CR) that exhibited rightward shifts in cisplatin IC50 (1.28-fold in HeLa and 1.80-fold in SiHa) and stable resistant phenotypes after drug withdrawal and cryopreservation. The present work therefore focuses on methodological standardisation of cisplatin resistance induction rather than exhaustive molecular characterisation, providing a clear starting point for future mechanistic and translational studies.

## 1 Introduction

Cervical cancer remains a significant global health burden, ranking as the fourth most frequently diagnosed malignancy and the fourth leading cause of cancer-related mortality among women worldwide [1].

In 2022 alone, an estimated 604,000 new cases and 342,000 deaths were reported globally annually, with disproportionately high incidence and mortality observed in low- and middle-income countries such as India [1]. These disparities largely reflect limited access to screening programmes, HPV vaccination, and timely treatment.

Cisplatin-based chemoradiation remains the standard first-line treatment for advanced cervical cancer [2]. By forming DNA–platinum adducts, cisplatin induces replication stress and apoptosis, leading to effective tumour regression in initially responsive cases. However, many patients eventually relapse as tumours acquire resistance to cisplatin, leading to treatment failure and disease recurrence [3].

In HPV-driven cervical cancer, viral oncoproteins can dysregulate apoptotic signalling and DNA repair mechanisms, facilitating a resistant cellular phenotype that undermines therapeutic outcomes [4].

Despite decades of research, the molecular underpinnings of cisplatin resistance remain incompletely defined, partly due to the lack of reproducible in vitro models. Traditional protocols for generating resistant cervical cancer cell lines vary widely in dose-escalation schedules, exposure durations, and validation criteria, often requiring months to achieve inconsistent results [5–7]. This methodological heterogeneity limits cross-study comparability and weakens mechanistic and preclinical conclusions regarding resistance biology.

To overcome these limitations, this study establishes a structured, stepwise protocol for developing cisplatin-resistant cervical cancer cell lines using HeLa (HPV18^+^) and SiHa (HPV16^+^) models. By incorporating fixed incremental dosing, systematic morphological monitoring, and half-maximal inhibitory concentration (IC_50_) recalculations, this approach enhances reproducibility and precision while reducing induction time. The resulting resistant lines were validated through comparative IC_50_ analyses and phenotypic assessment.

This work provides a scalable framework for generating resistance models and a translationally relevant platform for dissecting the molecular mechanisms driving cisplatin resistance in cervical cancer, thereby facilitating more reliable therapeutic screening and accelerating the identification of molecular targets for overcoming chemoresistance.

## 2 Rationale

Conventional protocols for developing cisplatin-resistant cervical cancer cell lines typically employ gradual dose-escalation, exposing parental cells to increasing concentrations of cisplatin across multiple passages [4, 8]. Despite the widespread use of these approaches, there is substantial methodological heterogeneity regarding dose increments, passage criteria, timing, and endpoints.

This has led to significant inter-laboratory variability, with induction timelines ranging from two to six months, fundamentally undermining the reproducibility and comparability of both mechanistic studies and preclinical drug screens across literature [4, 9]. In addition, existing methods often lack rigorous and standardised monitoring. Serial reassessment of drug uptake is inconsistently performed, while morphological surveillance is usually informal, overlooking subtle adaptive changes or non-specific stress responses [4, 8]. As a result, many reported resistant sublines risk accumulating artefactual phenotypes or genetic drift, diminishing their translational relevance as clinical resistance models.

To address these limitations, this study establishes a defined, stepwise induction protocol that integrates fixed 3 *µ*M increments of cisplatin with a minimum of three passages per dose, ensuring stable phenotypic adaptation before escalation. The protocol incorporates systematic, longitudinal monitoring of morphology and viability, coupled with IC_50_ recalculation at critical checkpoints. These built-in quality controls enable early detection of aberrant adaptation and enhance overall reproducibility.

The successful generation and validation of HeLa- and SiHa-derived resistant sublines (H_1_CR and S_1_CR), confirmed by statistically significant rightward IC_50_ shifts and stable post-treatment morphology, demonstrate that this approach produces robust and reproducible models of cisplatin resistance.

## 3 Results

Parental HeLa (RRID: CVCL 0030)[10] and SiHa (RRID: CVCL 0032)[11] cells were designated H_1_ and S_1_, respectively, and their resistant counterparts H_1_CR and S_1_CR.

### 3.1 Baseline IC_50_

Baseline IC_50_ values were calculated to establish the intrinsic cisplatin sensitivity of the parental cell lines. H_1_ and S_1_ cells showed IC_50_ values of **17.56** ±**0.85***µ*M and **14.71**±**0.89** *µ*M, respectively(mean ± SD, n=3), which were rounded to *18µ*M and *15µ*M for experimental purposes. These values served as reference points for subsequent resistance development and provided a quantitative baseline for assessing changes in drug responsiveness.

### 3.2 Development of cisplatin-resistant cell lines

HeLa and SiHa cells were subjected to the incremental cisplatin exposure protocol, resulting in the successful generation of the resistant sublines H_1_CR and S_1_CR. By the third passage at each concentration increment, cells demonstrated stable growth and regained proliferative capacity despite initial cytotoxicity.

#### 3.2.1 Morphological observations

Early passages showed characteristic stress responses, including partial detachment and cellular rounding. In later passages, both HeLa and SiHa cells exhibited normal adherent morphology (Fig. 1) and proliferation comparable to that of the parental lines. No significant morphological abnormalities were observed post-stabilisation.

**Fig. 1:**
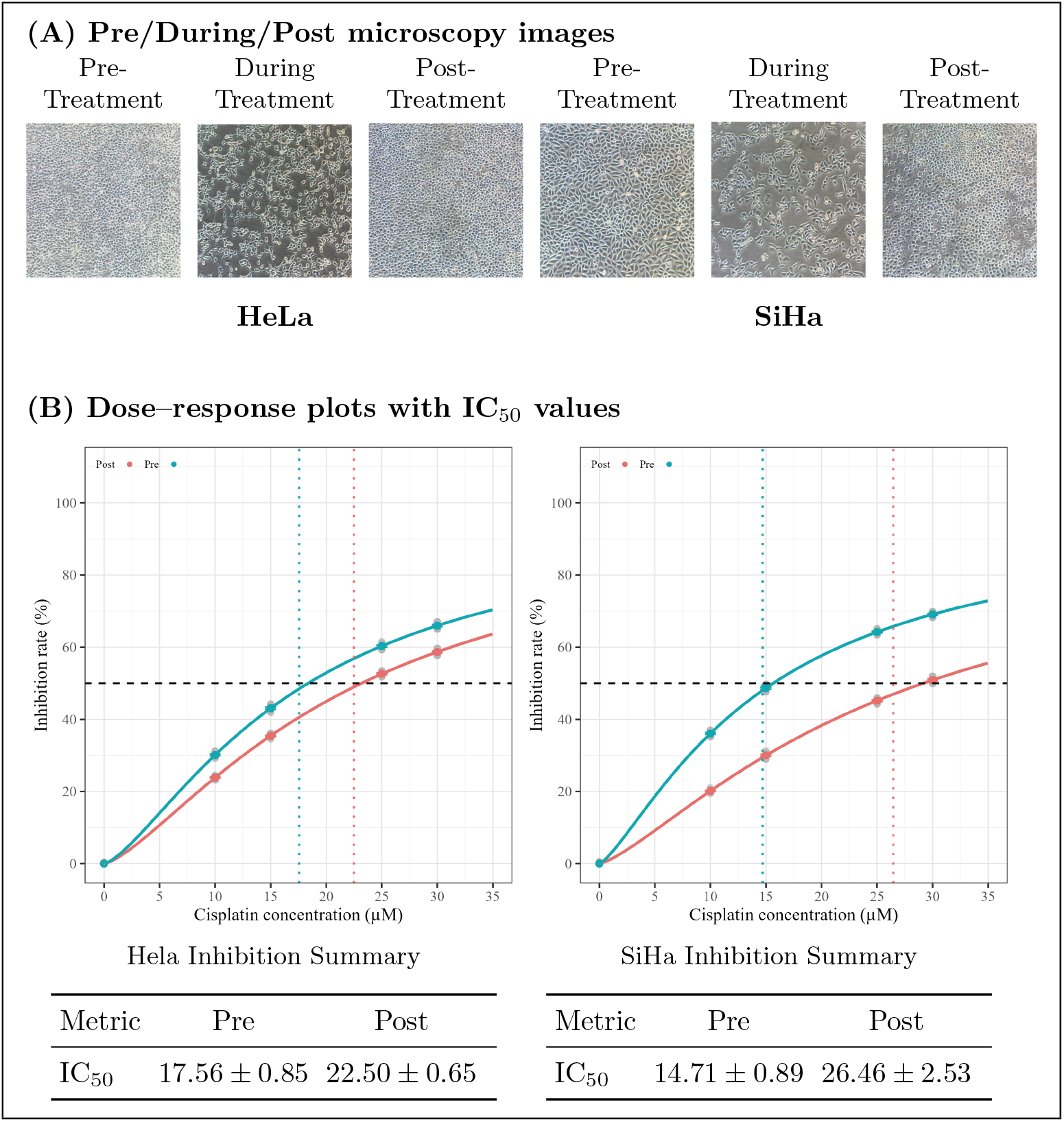
**(A)**: Representative phase-contrast images of HeLa and SiHa cells at pretreatment, during treatment, and post-treatment. **(B)** Dose–response curves for Pre vs Post sublines with IC_50_ indicated at 50% inhibition. Post-treatment IC_50_ values increased **1.28-fold in HeLa, p** = **0.0033** and **1.80-fold in SiHa, p** = **0.0018** (Two-tailed Welch’s t-test, *n* = 3 biological replicates); per-plot tables list IC_50_ means ± SD.

#### 3.2.2 Post-treatment IC_50_ values

Post-treatment IC_50_ values increased from 17.56 *µ*M to **22.50**± **0.65** *µ*M in HeLa (1.28 fold-change; *p* = 0.0033) and from 14.71 *µ*M to **26.46**± **2.53** *µ*M in SiHa (1.80 fold-change; *p* = 0.0018) (Fig. 1). These shifts, modest in HeLa and substantial in SiHa, confirm the stability of cisplatin resistance.

The elevated IC_50_ values were consistent across biological replicates, demonstrating that the induced resistance was stable immediately after the stepwise cisplatin exposure protocol and was not attributable to transient stress adaptation.

#### 3.2.3 Post-treatment cell health

Both H_1_CR and S_1_CR cell lines maintained a stable resistant phenotype after drug withdrawal. Following three passages in cisplatin-free medium, the resistant cells demonstrated rapid proliferative recovery, consistently reaching ∼90% confluence within 48 hours with no evidence of morphological stress, apoptosis, or growth retardation. Dose–response profiling confirmed that the elevated IC_50_ values were maintained, demonstrating that resistance was stable and not dependent on continuous drug pressure.

Resistance also persisted following cryopreservation. Cells revived from liquid nitrogen achieved ∼80% confluence within 36 hours post-thaw and displayed normal adherent morphology and expansion thereafter. IC_50_ values measured after thawing remained comparable to pre-preservation levels, indicating that cisplatin resistance was preserved through freeze–thaw cycling. Together, these observations confirm that the resistance phenotype is stable, heritable, and intrinsic to the selected cell populations.

## 4 Methods

### 4.1 Cell culture and maintenance

For protocol development, HeLa (HPV 18+ve cervical adenocarcinoma) and SiHa (HPV 16+ve cervical squamous cell carcinoma) were used. Both cell lines were maintained as adherent monolayer cultures using Dulbecco’s Modified Eagle Medium (DMEM) {HiMedia} supplemented with 10% Foetal Bovine Serum (FBS) {Gibco} and 1% Penicillin-Streptomycin antibiotic solution Thermo Fisher . Cells were routinely passaged at 90% confluence using 0.25% Trypsin-EDTA {Gibco}, to ensure optimal growth conditions.

### 4.2 Baseline IC_50_ determination

Before designing a dosing schedule, IC_50_ concentrations were determined using an Alamar Blue assay to establish each cell line’s initial drug sensitivity [12]. Determining baseline IC_50_ values and cisplatin uptake profiles provides a quantitative foundation for comparison with post-resistance cells, enabling more accurate assessment of treatment-induced changes. An initial broad-range dose–response assay was conducted to identify the appropriate experimental window, followed by a refined IC_50_ determination.

96-well plates were seeded with 10, 000 cells per well and incubated for 24 hours. The medium was then replaced with fresh culture medium containing cisplatin at concentrations of 10, 15, 20, 25 and 30*µ*M, along with a drug-free control. Following 24 and 48 hour exposure, 10*µ*L of Alamar Blue *{*Thermo Fisher*}* was added to each well, and fluorescence intensity was measured after a 3-hour incubation period using a Tecan Plate Reader.

Establishing these baseline IC_50_ values before resistance induction provides a consistent, data-driven framework for subsequent drug exposure protocols, enhancing methodological precision and improving comparability between parental and resistant cell lines.

### 4.3 Development of cisplatin-resistant cell lines

To establish cisplatin-resisnt cervical cancer models, the following exposure protocol was adopted.

#### 4.3.1 Stepwise exposure strategy

- Initial cisplatin exposure began at a common start of 13 *µ*M, with a 3 *µ*M incremental increase per cycle up to 30 *µ*M.
- Each exposure cycle consisted of 48 hours in cisplatin-containing medium, followed by recovery in drug-free medium until full confluence was regained (∼36-48 hours).
- Three passages per dose were performed to ensure stable phenotypic adaptation before increasing concentration.
- Throughout each cycle, morphological integrity, adherence, and viability were closely monitored via phase-contrast microscopy to detect early signs of cytotoxicity or protocol deviation.

#### 4.3.2 Cell recovery and stabilisation

- Despite initial cytotoxic stress, surviving clones gradually re-established proliferation under continued selection pressure.
- Following completion of dose-escalation, both H_1_CR and S_1_CR were cultured in drug-free medium for two weeks to stabilise their phenotype and mitigate transient stress responses.

This stepwise protocol, coupled with continuous morphological and viability assessment, minimised variability between replicates and enhanced reproducibility. The resulting H_1_CR and S_1_CR cell lines serve as stable models for investigating the cellular and molecular mechanisms underlying cisplatin resistance in cervical cancer.

A more detailed look into dosing and monitoring schedules can be found in Appendix 2.

### 4.4 Validation of cisplatin resistance

To confirm the successful establishment of cisplatin-resistant sublines (H_1_CR and S_1_CR), post-treatment IC_50_ values were recalculated using the same Alamar Blue dose–response assay described for the parental cell lines. Resistant cells were exposed to an adjusted concentration range (15–40 *µ*M) of cisplatin, and the resulting dose–response curves were compared to assess the degree of rightward shift in IC_50_ confirming reduced drug sensitivity and stable resistance acquisition.

To assess long-term stability, H_1_CR and S_1_CR cells were cryopreserved and revived before post-treatment analyses. Cells at near-confluence were frozen in FBS containing 10% DMSO and stored in liquid nitrogen for three weeks. Following revival, cells were cultured in drug-free medium under standard conditions (37°C, 5% CO_2_, humidified atmosphere) to allow recovery and expansion before downstream assays.

### 4.5 Data analysis

Raw fluorescence values from Alamar Blue assays were blank-subtracted and normalised to the mean signal of vehicle-treated control wells, which was set to 100% viability. Dose–response curves were then fitted for each experiment using a four-parameter log-logistic (LL.4) model implemented in the drc package in R. Statistical comparisons between parental and resistant lines were performed using two-tailed Welch’s t-tests (unpaired, unequal variances), with *p <* 0.05 considered statistically significant.

## 5 Discussion

The successful establishment of cisplatin-resistant HeLa (H_1_CR) and SiHa (S_1_CR) cell lines provides a stable model for investigating the molecular determinants of chemoresistance in cervical cancer. Resistance development, achieved through systematic incremental exposure and continuous phenotypic monitoring, yielded stable sublines that demonstrated significantly elevated IC_50_ values and morphological recovery consistent with adaptation rather than cytotoxic stress. These results confirm that the stepwise induction protocol effectively promotes gradual cellular adaptation while maintaining phenotypic continuity with the parental lines.

In contrast to previous conventional resistance-induction approaches [13, 14], this protocol standardises critical parameters including dose increments, passage thresholds, and IC_50_ recalculations, thereby reducing procedural variability. The increased IC_50_ values observed across both cell lines validate this approach and demonstrate that resistance development can be achieved efficiently, without prolonged selection or excessive cytotoxic attrition [12].

The morphological stability of H_1_CR and S_1_CR cells following two weeks of recovery in drug-free conditions suggests that resistance acquisition is intrinsic rather than transient [15]. Such stability is crucial for downstream applications, including geneexpression profiling and drug sensitivity screening. Incorporating quantitative IC_50_ reevaluation at multiple stages of selection and after drug-free recovery acts as a quality-assurance checkpoint to verify that the phenotype reflects stable resistance rather than transient stress tolerance. This approach aligns with recommendations to mitigate IC_50_’s timing and growth-rate dependencies and to pair it with robust dose–response practices [16].

This study has several limitations. First, the protocol was evaluated in only two cell lines, and the extent to which the same dose-escalation scheme and passage thresholds can be generalised to other cervical or non-cervical models remains to be determined. Although the observed increase in IC_50_ was substantial in SiHa, the fold change in HeLa was more modest, indicating that the dynamic range of resistance achievable with this protocol may be cell line–dependent. Second, all experiments were performed in two-dimensional monolayer culture, which does not fully recapitulate the threedimensional architecture, stromal interactions, or drug gradients present in vivo [17]. Finally, resistance was characterised primarily at the functional level using IC50 shifts and morphological recovery; comprehensive molecular profiling and long-term stability assessments will be required to define the resistant phenotype and its translational relevance fully.

To this end, further validation tests are being conducted on the established resistant lines to confirm their molecular and phenotypic stability. Ongoing analyses include short tandem repeat (STR) profiling to authenticate the genetic identity of H_1_CR and S_1_CR relative to their parental counterparts, ensuring that resistance was acquired without cross-contamination or genetic drift. In parallel, live-cell imaging studies are being performed to monitor dynamic cellular behaviours, such as proliferation, apoptosis, and morphological adaptation, under renewed cisplatin challenge.

Additionally, quantitative RT-PCR (qRT-PCR) assays are being conducted to profile differential expression of genes associated with DNA repair, apoptosis regulation, and drug efflux pathways, providing insight into the molecular mechanisms underpinning the resistant phenotype. These validation experiments will further strengthen the translational relevance of the established models [14, 18].

Taken together, these findings establish a framework for generating cisplatin-resistant cervical cancer models. Beyond serving as a methodological proof-of-concept, the protocol provides a reliable foundation for mechanistic studies and preclinical drug testing. By bridging experimental precision with translational applicability, this work advances the development of tools for dissecting resistance biology and accelerating therapeutic innovation in cervical cancer management.

## 6 Conclusion

This study establishes cisplatin-resistant HeLa (H_1_CR) and SiHa (S_1_CR) cell lines using a transparent, stepwise induction protocol with predefined dose increments, passage criteria, and IC_50_ checkpoints. The protocol reliably generates stable resistant sublines that retain normal adherent morphology and show reproducible rightward shifts in cisplatin dose–response curves.

By making each experimental decision point explicit, this framework reduces the ambiguity that has historically limited comparability between cisplatin-resistance models. Although currently only validated in two HPV-positive cervical cancer lines under two-dimensional culture, following further validation, the workflow can be readily adapted to other cell types, drug regimens, and more complex culture systems. Together, H_1_CR and S_1_CR provide an accessible platform for functional assays, molecular profiling, and preclinical testing of combination strategies to prevent or overcome cisplatin resistance in cervical cancer.

## 7 Acknowledgements

I acknowledge the Cancer Institute (WIA), Research Wing, Department of Molecular Oncology, for providing the infrastructure for this research, and thank Dr R. K. Sabitha and Dr Deva Magendhra Rao A. K for supervision, Hema and Deepa for laboratory training and support, the research staff, particularly Muthu, for administrative assistance, and Dyuthi, Sasi, Daniel and Ananth for help with troubleshooting experiments and data analysis.

## A Detailed Resistance Development Protocol

### A.1 Experiment Setup

Protocol performed on HeLa (HPV 18+ve cervical adenocarcinoma) and SiHa (HPV 16+ve squamous cell carcinoma) cell lines.

- Cells were maintained in Dulbecco’s Modified Eagle Medium (DMEM) supplemented with 10% Foetal Bovine Serum (FBS) and 1% Pencillin-Streptomycin antibiotic solution.
- Cells were grown as monolayer cultures at 37°in a humidified incubater with 5% CO_2_.

### A.2 Cisplatin Exposure

After initial IC_50_s are obtained, define the shared starting dose as the lower of the two IC_50_ values.

1. Seed cells in T25 flasks to reach ∼100% confluence after 24h
2. Prepare a 1 mg/mL cisplatin stock in 0.9% NaCl in a light-protected tube, then dilute into prewarmed medium to the desired working concentrations immediately before use.
3. Aspirate spent medium and add fresh, prewarmed complete medium with cisplatin at the determined dose.
4. Incubate cells in cisplatin-containing medium for 48 h, replacing the medium at 24 h with fresh cisplatin-containing medium to maintain drug concentration.
5. After 48 h, remove the cisplatin-containing medium and wash cells twice with 2 mL Ca^2+^/Mg^2+^-free PBS (room temp)
6. Add trypsin to detach cells; neutralize with 3 mL prewarmed complete medium and transfer to a 15 mL tube.
7. Centrifuge at 300 × g (≈1000 rpm) for 3 min; discard the supernatant.
8. Resuspend the cell pellet in drug-free complete medium and reseed into fresh T25 flasks at the desired density.
9. Once cells recover and reach 80–90% confluence, subculture into fresh T25 flasks.
10. Re-expose for 48 h, then culture in drug-free medium until confluence is reestablished.
11. Repeat this stepwise exposure for 3 passages at each dose increment to promote stable adaptation.
12. Increase cisplatin concentration by 3*µ*M each cycle.

### A.3 Monitoring and Quality Control

1. **Cell morphology**: Examine cultures on an inverted microscope for changes in shape, granularity, and spontaneous detachment.
2. **Viability assay (Trypan Blue)**: Mix a cell aliquot 1:1 with 0.4% Trypan Blue and count live/dead cells using a hemocytometer or automated counter.
3. Alamar Blue assay:
4. Add 10*µ*L Alamar Blue per well in a 96-well plate (to 100*µ*L medium).
5. Incubate for ∼3 h at 37°C, 5% CO_2_.
6. Read fluorescence (Ex ∼560 nm/Em ∼590 nm) or absorbance (570/600 nm), blank-subtracted.

### A.4 Achieving Stable Resistance

1. Continue stepwise cisplatin exposure until the target resistance level is reached (per predefined IC_50_ or fold-change criterion).
2. Maintain cells in drug-free medium for two weeks to stabilise the phenotype.
3. Perform routine medium changes every 48 h.

### A.5 Cryopreservation Of Resistant Clones

1. When cultures reach 80–90% confluence, detach with trypsin and collect cells.
2. Centrifuge at ∼300 × g (≈1000 rpm; rotor-dependent) for 3 min and discard the supernatant.
3. Resuspend the pellet in ice-cold freezing medium (90% FBS + 10% DMSO) at ∼0.5™ 1.9* 10^6^ cells/mL.
4. Aliquot 1 mL per cryovial and cool at ∼1 °C/min to -80°C (e.g., isopropanol freezing container) overnight.
5. Transfer vials to liquid nitrogen for long-term storage.

## References

[1] Bray, F. et al. Global cancer statistics 2022: GLOBOCAN estimates of incidence and mortality worldwide for 36 cancers in 185 countries. CA: A Cancer Journal for Clinicians 74, 229–263 (2024).

[2] Yang, L. et al. Molecular mechanisms of platinum-based chemotherapy resistance in ovarian cancer (Review). Oncology Reports 47, 1–11 (2022).

[3] Wang, L., Zhao, X., Fu, J., Xu, W. & Yuan, J. The Role of Tumour Metabolism in Cisplatin Resistance. Frontiers in Molecular Biosciences 8 (2021).

[4] Zhu, H. et al. Molecular mechanisms of cisplatin resistance in cervical cancer. Drug Design, Development and Therapy 10, 1885 (2016).

[5] Brown, A., Kumar, S. & Tchounwou, P. B. Cisplatin-Based Chemotherapy of Human Cancers. Journal of cancer science & therapy 11, 97 (2019).

[6] Bahar, E., Kim, J.-Y., Kim, H.-S. & Yoon, H. Establishment of Acquired Cisplatin Resistance in Ovarian Cancer Cell Lines Characterized by Enriched Metastatic Properties with Increased Twist Expression. International Journal of Molecular Sciences 21, 7613 (2020).

[7] Shirmanova, M. V. et al. Chemotherapy with cisplatin: Insights into intracellular pH and metabolic landscape of cancer cells in vitro and in vivo. Scientific Reports 7, 8911 (2017).

[8] Bhattacharjee, R. et al. Cellular landscaping of cisplatin resistance in cervical cancer. Biomedicine & Pharmacotherapy 153, 113345 (2022).

[9] Galluzzi, L. et al. Systems biology of cisplatin resistance: Past, present and future. Cell Death & Disease 5, e1257 (2014).

[10] CALIPHO group, SIB Swiss Institute of Bioinformatics. Cellosaurus: Hela (rrid: Cvcl 0030) (2025). URL https://www.cellosaurus.org/CVCL0030. Database record; Resource Identification Initiative.

[11] CALIPHO group, SIB Swiss Institute of Bioinformatics. Cellosaurus: Siha (rrid: Cvcl 0032) (2025). URL https://www.cellosaurus.org/CVCL0032. Database record; Resource Identification Initiative.

[12] Bonnier, F. et al. Cell viability assessment using the Alamar blue assay: A comparison of 2D and 3D cell culture models. Toxicology in vitro: an international journal published in association with BIBRA 29, 124–131 (2015).

[13] Mahapatra, E. et al. in Insights of Cisplatin Resistance in Cervical Cancer: A Decision Making for Cellular Survival (ed. Rajkumar, R.) Cervical Cancer - A Global Public Health Treatise (IntechOpen, 2021).

[14] Roy, M. & Mukherjee, S. Reversal of Resistance towards Cisplatin by Curcumin in Cervical Cancer Cells. Asian Pacific Journal of Cancer Prevention 15, 1403–1410 (2014).

[15] Nunes, M. et al. Generation of Two Paclitaxel-Resistant High-Grade Serous Carcinoma Cell Lines With Increased Expression of P-Glycoprotein. Frontiers in Oncology 11 (2021).

[16] Hafner, M., Niepel, M., Chung, M. & Sorger, P. K. Growth rate inhibition metrics correct for confounders in measuring sensitivity to cancer drugs. Nature Methods 13, 521–527 (2016).

[17] Fang, Y. & Eglen, R. M. Three-Dimensional Cell Cultures in Drug Discovery and Development. SLAS Discovery 22, 456–472 (2017).

[18] Huang, D. et al. A highly annotated database of genes associated with platinum resistance in cancer. Oncogene 40, 6395–6405 (2021).

